# The forbidden doubling: exploring rare spermatocyte polyploidy in mammals

**DOI:** 10.1101/2025.10.25.684494

**Authors:** Sergey Matveevsky, Oxana Kolomiets, Tatiana Grishaeva, Aleksey Bogdanov, Valentina Tambovtseva, Irina Bakloushinskaya

## Abstract

We studied rare cases of over-ploid spermatocytes using advanced immunocytochemical methods and a cross-species approach for the first time. In subterranean rodents *Ellobius tancrei, E. alaicus, E. talpinus*, and *Nannospalax leucodon* tetraploid spermatocytes exhibited specific features during meiotic prophase I, including symmetric and asymmetric chromosome quadrivalents with partner-switching, extended asynapsis, altered recombination patterns, and variable chromatin inactivation. Notably, the quadrivalents were capable of assembling shelterin complexes at chromosome ends, which are connected to the nuclear envelope through the SUN–KASH complex as in normal spermatocytes. Anomalies suggest that meiotic checkpoints potentially triggered by failed synapsis or incomplete sex chromosome silencing may act to prevent the progression of polyploid spermatocytes.

## Introduction

Polyploidy, i.e. the presence of more than two complete chromosome sets in some or all cells of an organism, is widespread among eukaryotes and contributes to evolutionary diversification, particularly in invertebrates and lower vertebrates. Polyploidy in somatic cells in liver, pancreas, heart, and muscles, was well documented in animals and humans (Fox and Duronio 2013; Wang et al. 2017). Certain tissues in females of mammals (in the placenta and lactating mammary glands) also exhibit polyploidy, which may enhance the organ function (Zybina and Zybina 2020; Anatskaya and Vinogradov 2022).

Despite numerous cases of somatic polyploidy, several factors constrain the whole-genome duplication in mammals. The presence of the heteromorphic sex chromosomes was hypothesized as the most affective to prevent polyploidy establishing (Ohno 1970; Stöck et al. 2021). In addition, the increased nuclear volume and chromosome number in polyploid cells may compromise nuclear architecture, interfere with chromosome interactions, and impair mitotic segregation (Shu et al. 2018; Vittoria et al. 2023). Finally, polyploidy is exceptionally rare in the germline in mammals and is viewed as a deviation from haploid gamete production. This is due to the fact that polyploidy in gametogenesis can potentially produce aneuploid progeny (Mason and Pires 2015). Furthermore, although diploid gametes may be formed through abnormal mitotic divisions, such as endoreduplication or failed cytokinesis (Therman and Susman 1993; Egozcue et al. 2000; Escalier 2002; Lee et al. 2009; Zielke et al. 2013; Stenberg and Saura 2013), these cells are typically eliminated during meiotic checkpoints, especially in males (Li et al. 2009). In contrast, female meiosis appears to be more error-prone, and diploid oocytes are more likely to contribute to over-ploid embryos in humans (Beatty and Fischberg 1952; Speed 1985; Capalbo et al. 2017; Samura et al. 2023).

There are only a few reports of polyploid spermatocytes in model mammalian organisms (Lin et al. 1971; Pogosianz and Brujako 1971; Chandley et al. 1974; Holm and Rasmussen 1983) and humans (Carothers and Beatty 1975; Kajii and Niikawa 1977; Wegner et al. 2001; Sarrate et al. 2014; Pearson and Madan 2018). Although sporadic instances of meiotic polyploidy have been reported in mammals, cytologically characterized cases of autotetraploid spermatocytes, particularly those with fully assembled synaptonemal complexes (SCs), remain exceedingly rare. To date, only isolated cases have been reported in mice (Solari and Moses 1977), humans (Codina-Pascual et al. 2006; Schmidt 2017), northern mole vole (Matveevsky et al. 2017), and recently in brown bears (Virabyan et al. 2025), leaving major gaps in our understanding of how polyploid germ cells behave during meiosis and how they are regulated. The mechanisms by which tetraploid meiocytes progress through early prophase I, resolve complex multivalent configurations, and interact with meiotic surveillance systems remain largely unknown.

In this study, we aimed to identify and characterize rare tetraploid spermatocytes in selected rodent species, including mole voles and mole rats, using immunocytochemistry. We focused on the structural configurations of the SCs, recombination activity, and chromatin inactivation dynamics during meiotic prophase I. By comparing these features with those observed in diploid cells, we sought to provide cytological insight into the fate of naturally occurring polyploid spermatocytes and to explore the potential involvement of meiotic checkpoints that may eliminate or arrest cells carrying unresolved multivalents. By integrating comparative cytogenetic data with functional markers of telomere attachment (RAP1) and nuclear envelope interactions (SUN1), our study elucidates how mammalian meiosis restricts polyploid germ cell formation and maintains genomic stability.

## Materials and methods

In total, 11 polyploid spermatocytes were studied in five males of four rodent species. *Nannospalax leucodon* (the mole rat) was captured near Ravno Pole village, eastern Sofia, Bulgaria (Matveevsky et al. 2021). Two males of the Eastern mole vole *Ellobius tancrei* were obtained in prolonged experimental breeding (2n = 49, ID #25187 and 2n = 49, ID #27430, see [Kolomiets et al. 2023]). *Ellobius alaicus* (the Alay mole vole) male (2n = 52, ID #27495) from southeastern Kyrgyzstan and *E. talpinus* (the common mole vole) male (2n = 54, ID #27416), the offspring of experimental breeding, were provided by the Large-Scale Research Facility “Collection of Wildlife Tissues for Genetic Research,” IDB RAS, registration no. 3579666. All procedures followed institutional and international ethical standards (Ethics Committee approvals: VIGG RAS Order No. 3, 10 November 2016; IDB RAS Protocol No. 37, 25 June 2020).

Preparations of meiotic chromosomes at the stage of diakinesis and metaphase I regularly reveal cells, resembling tetraploid ones, but it is difficult to prove that the cells are true polyploid because of their possible overlapping. The best way to prove genuine tetraploidy of spermatocytes is to identify nuclei in the middle of prophase I, when chromosome synapsis occurs. Cell spreads for studying synaptonemal complex (SC) were prepared following Peters et al. (1997) (some details see Gil-Fernández et al. 2021).

The following primary antibodies were used: anti-SYCP3 (rabbit, 1:250, Abcam, UK) to label axial/lateral elements of SCs; anti-centromere (CREST) (human, 1:250, Fitzgerald, USA) to identify centromeres; anti-γH2AFX (mouse, 1:250–500, Abcam) to detect chromatin inactivation; anti-MLH1 (mouse, 1:50, Abcam) as a recombination marker; anti-RAP1 (rabbit, 1:100, Abcam) to detect telomere sites; Anti-SUN1 (rabbit, 1:250, Abcam) to visualize nuclear envelope LINC components. Secondary antibodies included goat anti-rabbit IgG Alexa Fluor 488, anti-human IgG Alexa Fluor 546, and anti-mouse IgG Alexa Fluor 546/555 (Invitrogen, USA; dilution 1:300–800). Slides were washed in PBS and mounted with Vectashield containing DAPI (Vector Laboratories, USA). Visualization was performed using an Axio Imager D1 fluorescence microscope (Carl Zeiss, Germany). The immunostaining protocols followed previously described procedures (Matveevsky et al. 2021).

A “quadrivalent” designation was used in this paper to characterize SC configurations consisting of 4 axes in tetraploid meiocytes, in contrast to a “tetravalent” usually used when describing SC configurations in chromosomally heterozygotic (hybrid) organisms, such as in our previously published work (Matveevsky et al. 2015).

## Results and discussion

Meiotic spreads from *E. tancrei, E. alaicus, E. talpinus*, and *N. leucodon* revealed several tetraploid spermatocytes. These were confidently distinguished from overlapping diploid cells by analyzing synapsis patterns at prophase I and using multiple immunocytochemical markers.

### Case Study 1: *Ellobius tancrei*

Among thousands of spermatocytes examined in chromosomally variable *E. tancrei*, only two cells were clearly tetraploid (Figs 1, S1). #27430, 2n=49 specimen was heterozygous for one Robertsonian translocation and previously shown to be sterile (Kolomiets et al. 2023). The tetraploid cell exhibited four SC quadrivalents, two univalents, and several trivalent-like configurations. Four X chromosome axes were centrally located within a γH2AFX-positive chromatin cloud, with two fragmented Xs.

**Figure 1.**
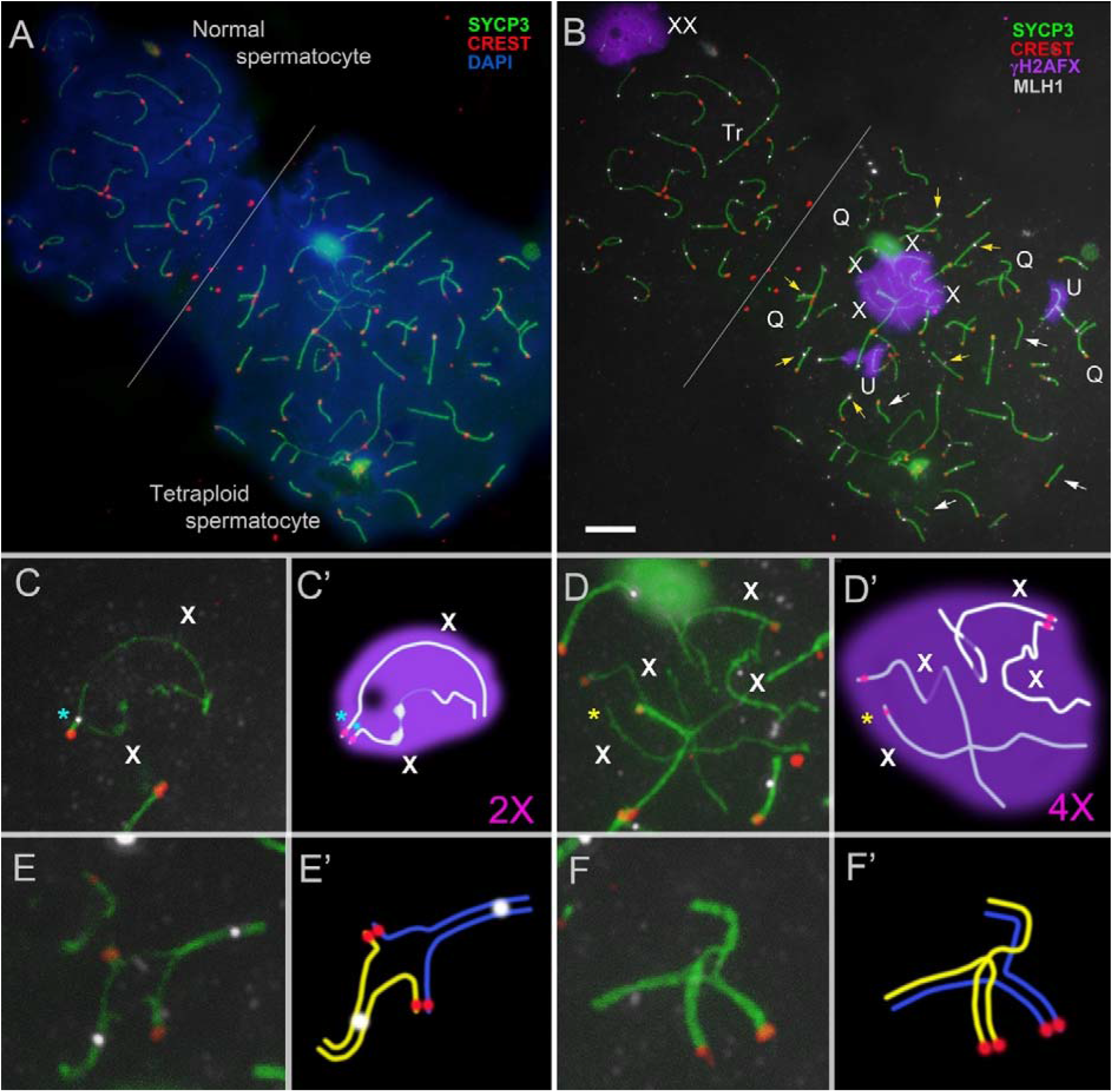
Normal (diploid) and tetraploid pachytene spermatocytes of *E. tancrei* #27430, 2n=49. Axial and lateral elements of the SCs were identified using an anti-SYCP3 antibody (green); centromeres were identified using “CREST”—antibodies to kinetochore proteins (red); anti-MLH1 antibodies (white) were used as a marker of recombination nodules; and chromatin inactivation was revealed using an anti-γH2AFX antibody (violet). Chromatin was stained with DAPI (blue). X – sex chromosomes, Q – chromosomal quadrivalent, Tr – chromosomal trivalent, U – univalent. Both spermatocytes were photographed in one microscopic image without changes in their position. Normal and tetraploid spermatocytes differ from each other in cell size (see DAPI in A) and the presence of quadrivalents in the latter (see Q in B). White arrow points to SC bivalents without MLH1 dots (B). Yellow arrow indicates SC bivalents with shifted MLH1 foci (B). A blue asterisk indicates the presence of a recombination nodule in the sex (2X) bivalent in the normal spermatocyte (C, C’). In a tetraploid spermatocyte, four axes were observed within one γH2AFX cloud, which corresponded to four X chromosomes (4X) (B, D, D’). The centromeric region is not visualized on one of the X axes (yellow asterisk). One quadrivalent has two MLH1 dots (E, E’), another quadrivalent is without MLH1 signals (F, F’). Scale bar = 5 μm.

MLH1 analysis revealed 39 recombination foci in the tetraploid cell that is higher than the diploid average (21.6) and supports its tetraploid status. Some bivalents lacked MLH1 foci, while others displayed shifted or atypically located foci.

A single tetraploid diplotene cell was found in male #25187, 2n=49 (Fig. S1). The presence of multivalents due to Robertsonian translocations and univalents compromises the normal end of meiotic prophase I and probably leads to elimination of the cell.

### Case Study 2: *Ellobius alaicus*

Two tetraploid nuclei were identified in a specimen of *E. alaicus*. SC quadrivalents showed symmetrical configurations with asynapsis in proximal regions and single crossover switches (Figs S2, S3). SUN1 immunostaining revealed chromosomal end-attachment to the nuclear envelope, implicating LINC complex involvement in SC architecture. Sex chromosomes were not recognized because of insufficient morphological and immunocytochemical specificity.

### Case Study 3: *Ellobius talpinus*

In a tetraploid spermatocyte of the common mole vole *E. talpinus*, both symmetric and asymmetric quadrivalents with and without partner switching were visible (Fig. S4A-L). We previously found similar quadrivalents in this species (Matveevsky et al. 2017). Immunodetection of RAP1 protein revealed for the first time that both quadrivalents and resolved bivalents (escaped from their quadrivalent configurations) formed shelterin complexes in telomeric regions (Fig. S4C-L). Of particular interest is the finding that both γH2AFX-positive and γH2AFX-negative axes were identified as a part of the quadrivalents in this cell (see white and cyan arrows in Fig. S4B). Sex chromosomes could not be reliably distinguished due to lack of morphological and immunocytochemical markers.

### Case Study 4: *Nannospalax leucodon*

In the 56-chromosomal form of *N. leucodon*, six tetraploid spermatocytes were found among 251 cells at different pachytene substages (Fig. S5). In the presented tetraploid spermatocytes of the mole rat, the number of centromeres was approximately 2-fold higher (58 instead of 28 in diploids), supporting tetraploidy. Chromosomes appeared as SC bivalents and various quadrivalent-like formations. We were unable to identify sex chromosomes.

Quadrivalents displayed both symmetric (fully synapsed arms) and asymmetric configurations (incomplete synapsis or desynapsis). Partner-switching within quadrivalents was common, and asynapsis was prominent in certain regions.

The analysis of four rodent species revealed that tetraploid spermatocytes occur rarely and display consistent cytological features. These abnormal cells were characterized by the formation of quadrivalents with partial asynapsis, elevated numbers of MLH1 recombination foci, and distinct abnormalities of sex chromosome including their central localization, fragmentation, and intensive γH2AFX staining. Additionally, the SUN1-positive chromosomal ends indicated nuclear envelope anchoring of quadrivalents. Together, these findings demonstrate that while tetraploidy in male rodent meiosis occurs sporadically, it produces structurally unique meiotic configurations that differ fundamentally from normal diploid spermatocytes.

Constitutional polyploidy is characteristic for a large number of plants and lower vertebrates, but there are no such examples for mammals (Otto and Whitton 2000; Gregory and Mable 2005; Song et al. 2012). In *Tympanoctomys barrerae*, GISH (genomic in situ hybridization) on mitotic metaphase and meiotic (the pachytene stage) cells and chromosome painting support allotetraploidy (Gallardo et al. 1999; Suárez-Villota et al. 2012), but this remains a unique case.

Since the discovery of chromosomes, studies of gametogenesis in polyploid organisms, particularly meiosis, have been in a focus of attention (Newton and Darlington 1929; White 1933; McCollum 1957). In this context, the processes of synapsis and chromosome recombination in prophase I of meiosis have been mainly emphasized in plants and sometimes insects (Rasmussen 1986; Higgins et al. 2014). Despite the fact that cases of sporadic meiotic polyploidy have been identified, autotetraploid meiocytes with SCs are detected very rarely. To our knowledge, to date, if we include the results of this work, there are only 9 cases of studying SCs in tetraploid spermatocytes: 2 in humans, 1 in bear and 6 in different rodent species (Table 1).

**Table 1.**
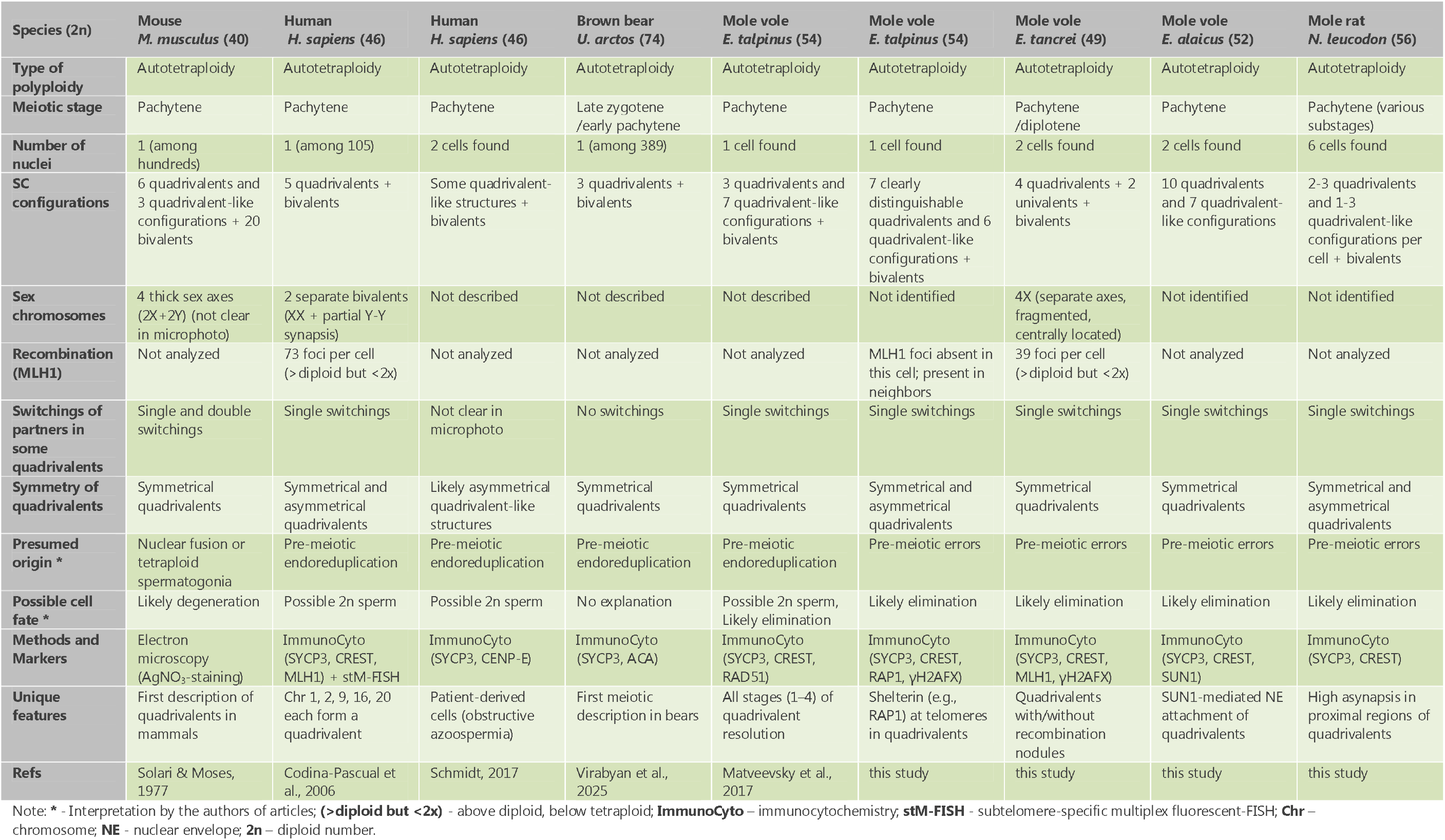
Synaptonemal complexes in mammalian tetraploid spermatocytes.

This study provides rare cytological evidence of polyploid meiosis in four rodent species and reveals consistent anomalies during prophase I. The most prominent feature across all species was disturbed synapsis, particularly in sex chromosomes, which likely disrupts progression through meiotic checkpoints and prevents the next generation of viable diploid sperm.

Our observations (here and Matveevsky et al. 2017) and simulations suggest a dynamic sequence of transitions from quadrivalents to individual bivalents, involving a series of distinct stages. These transitions exemplify an adaptive synaptic strategy that enables sporadically tetraploid cells to restore diploid-like pairing configurations and preserve meiotic ability to progression. The resolution dynamics of quadrivalents during meiotic prophase I in sporadic tetraploid cells can be summarized through four distinct stages (Fig. 2).

**Figure 2.**
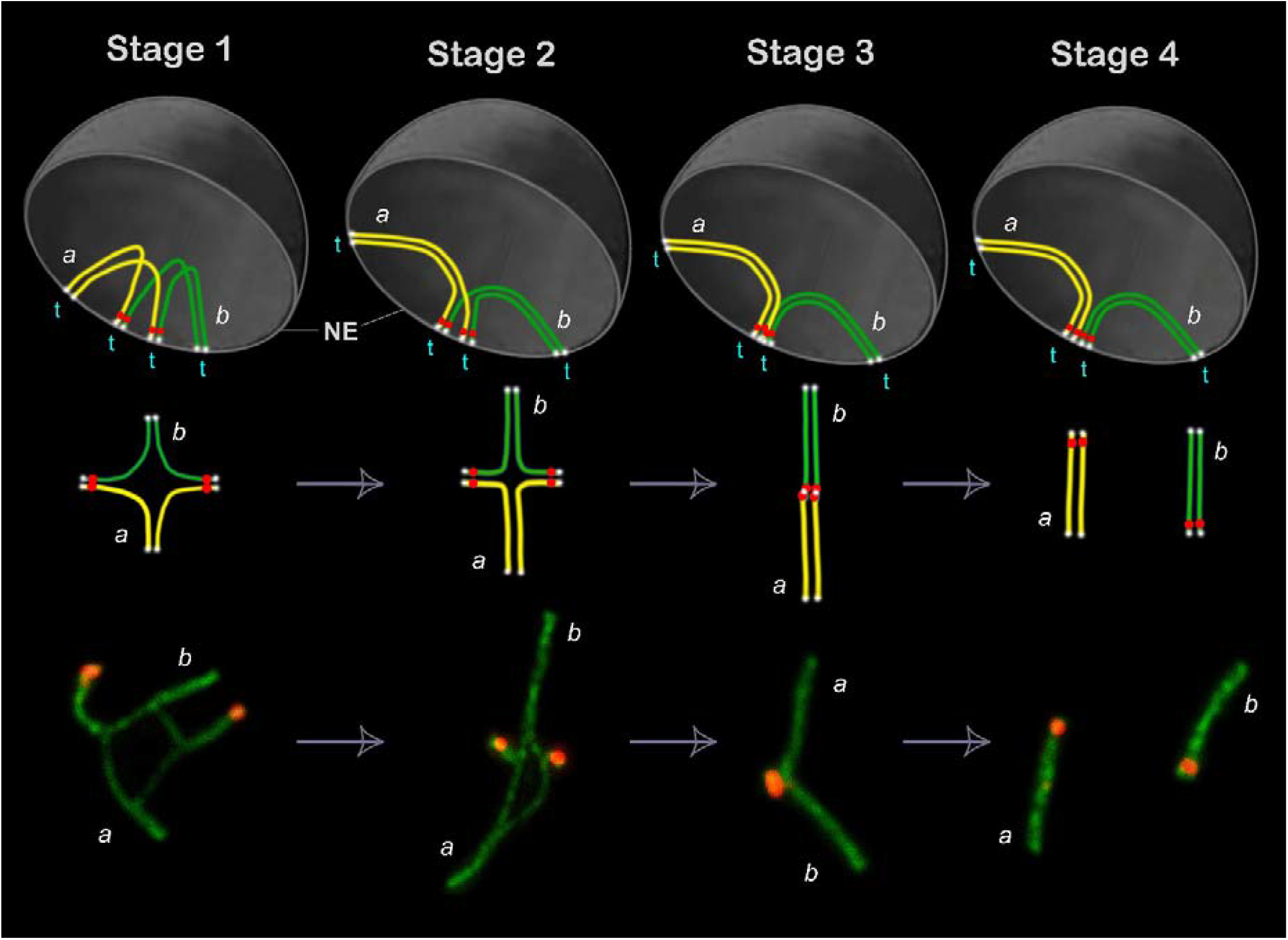
Synaptic adjustment and multivalent resolution: modeling the transition from quadrivalents to bivalents during meiotic sporadic tetraploidy. Top row – simulation of quadrivalents within meiotic nuclei. Middle row – schematic representation of quadrivalent dynamics. Bottom row – examples of quadrivalent and bivalent configurations derived from a single cell (immunocytochemistry: axial and lateral elements of the SCs are shown in green, centromeres in red). Abbreviations: NE – nuclear envelope; t – telomeres attached to the nuclear envelope (represented as white dots at chromosome ends in the top and middle panels). Color coding: a – yellow homologs; b – green homologs; red dots correspond to centromeres. Stage 1: Partial synapsis of four homologs (2 yellow, 2 green) forms a quadrivalent with sometimes broad asynaptic central region; configurations are often symmetrical. Stage 2: Synapsis progresses between homologous arms, reducing synaptic interactions between yellow and green homologs. Stage 3: Loss of synapsis between yellow and green homologs results in an extended, sometimes slightly curved, linear-like SC - resembling a bivalent, but unresolved; termed a pseudobivalent. Stage 4: Complete resolution of the quadrivalent into two bivalents.

The stage 1 was designated ‘Quadrivalent assembly through partial synapsis of four homologs’. This initial stage is characterized by partial synapsis among four homologous chromosomes (two yellow and two green in the model of nucleus, Fig. 2), leading to the establishment of a quadrivalent. Synapsis generally occurs along the arms of homologous chromosomes, frequently producing a symmetrical configuration. A hallmark of this stage is the presence of an extensive asynaptic region near the center of the quadrivalent, reflecting regions of misaligned or incomplete pairing. This apparent symmetry may reflect a relatively balanced mechanical tension exerted on the telomeric ends, which remain anchored to the nuclear envelope. Here, we demonstrate for the first time that quadrivalent chromosome configurations are capable of assembling fully formed shelterin complexes at telomeric ends, which are connected to the nuclear envelope through the SUN–KASH (LINC) complex, as in normal spermatocytes.

The stage 2 was designated ‘Synaptic refinement and reconfiguration of quadrivalent’. As meiosis progresses, synapsis becomes increasingly restricted to true homologous arms (i.e., yellow-yellow and green-green in Fig. 2), while synaptic interactions between heterologous homologs (yellow-green in Fig. 2) diminish. This results in a quadrivalent with two long and two short synapsed arms.

We designated the stage 3 as ‘Transitional linearization of a quadrivalent into a pseudobivalent’. At this stage, synapsis between heterologous homologs is almost entirely lost, giving rise to a long, continuous SC structure. This SC is often linear and can exhibit mild central curvature, likely due to telomere-nuclear envelope attachments. Despite its visual resemblance to a bivalent, this structure retains features of a multivalent origin and has not undergone full resolution. We therefore designate it a pseudobivalent.

The stage 4 is ‘Final quadrivalent resolution and formation of bivalents’. During the final stage, the quadrivalent structure fully resolves into two discrete bivalents, each formed by homologous pairing of the original chromosome sets (yellow-yellow and green-green in Fig. 2). The resulting bivalents are cytologically indistinguishable from those formed in normal meiocytes, indicating the successful correction of an initially aberrant tetraloid synaptic configuration.

Polyploid cells identified at the pachytene stage most likely originated due to errors during premeiotic mitoses. Although some resemble syncytial fusions (Figs S2, S3, and S5), others, like the cells in *E. tancrei* and *E. talpinus*, appear to be genuine tetraploids (Figs 1, S1, S4).

Several key findings were revealed in synaptic configuration, recombination events, and transcriptional inactivation of meiotic chromatin. Quadrivalents in tetraploid spermatocytes generally displayed symmetric synapsis in distal regions and asynapsis proximally. Some quadrivalents resembled adjacent bivalents connected at telomeric ends, suggesting partial resolution of quadrivalents; this process likely assisted by SUN1-mediated nuclear anchoring. Despite asynapsis, recombination foci (MLH1) were present in many quadrivalents and bivalents, albeit with altered positioning. This indicates that some homolog interactions and crossover formation remain functional, though possibly desynchronized or spatially misregulated.

Identification of sex chromosomes by γH2AFX staining revealed four unsynapsed axes in *E. tancrei* spermatocytes, which remained centrally located rather than migrating to the nuclear periphery as typically observed in normal spermatocytes (Fig. 1). These axes (univalents) were γH2AFX-positive, indicating asynapsis, whereas asynapsed regions within quadrivalents lacked γH2AFX signals, suggesting prior quick synapsis followed by desynapsis and possible evasion of meiotic silencing mechanisms (Burgoyne et al. 2009). A similar pattern was observed in *E. talpinus*, where quadrivalents exhibited both γH2AFX-positive and γH2AFX-negative axes. It is hypothesized that chromosome regions, lacking γH2AFX, have already undergone a brief synapsis phase followed by desynapsis. Meanwhile, γH2AFX-positive regions likely remain unsynapsed. This temporal desynchronization of synapsis may reflect genomic instability in tetraploid meiotic cells.

Asynapsis and improper recombination are well-known triggers of meiotic arrest (Turner et al. 2005; Subramanian and Hochwagen 2014). Our findings support the idea that disrupted synapsis and meiotic sex chromosome inactivation (MSCI) failure in tetraploid cells can activate checkpoint pathways, leading to meiotic arrest and likely elimination (Li et al. 2009). Rare escape from these checkpoints could explain the exceptionally low frequency of diploid spermatozoa reported in humans: from 0.0001% to 0.34% in different studies (Lu et al. 1994; Rodrigo et al. 2010).

Although rare polyploid embryos do occur, they are usually mosaics with severe abnormalities and early mortality (Cavalcanti DP, Zanchetta 2005; Schacht et al. 2018). Stable germline transmission of polyploid genomes in mammals remains virtually nonexistent.

The rare occurrence of polyploid spermatocytes in the four studied rodent species highlights polyploidy as a notable deviation in mammalian meiosis. Such cells consistently display disrupted synapsis, particularly for sex chromosomes, as well as aberrant recombination patterns and meiotic inactivation anomalies. Such abnormalities likely trigger early meiotic checkpoints and lead to cellular elimination before spermiogenesis.

Although tetraploid cells may occasionally proceed further through meiosis, the resulting spermatozoa are typically non-viable or eliminated, and their participation in fertilization is exceedingly rare. Cases of over-ploid embryos in humans are associated with severe developmental abnormalities and do not support heritable polyploidy.

Our findings reinforce the hypothesis that mammalian sex chromosomes, through mechanisms such as MSCI, serve as evolutionary safeguards against polyploid genome inheritance. So, polyploidy in mammalian gametogenesis is a genetic anomaly rather than an adaptive mechanism. Meiotic checkpoints and nuclear architecture prevent the inheritance of such overcomplete genomes, making polyploid evolution highly improbable in mammals, unlike in plants or lower vertebrates.

## Authors’ contributions

Conceptualization: S.M., O.K. and I.B.; methodology: O.K. and S.M.; investigation: S.M., T.G., V.T., A.B. and I.B.; writing: S.M., V.T., A.B., I.B. and O.K.; visualization: S.M. and I.B. All authors have read and agreed to the published version of the manuscript.

## Acknowledgments

This study is co-authored by Oxana Kolomiets, who sadly passed away before the submission of the manuscript. Her profound scientific insight, tireless dedication, and invaluable input shaped this and many other works. We deeply appreciate the years of collaboration, her mentorship, and her scientific legacy. We thank Sirma Zidarova, Nasko Atanasov, and Tsenka Chassovnikarova for their participation in the fieldwork and organizational support.

## Supplementary Information

The online version contains supplementary material available at…

