## Supplemental figures for "The forbidden doubling: exploring rare spermatocyte polyploidy in mammals"

* Professor Oxana Kolomiets died prior to the submission of this paper.

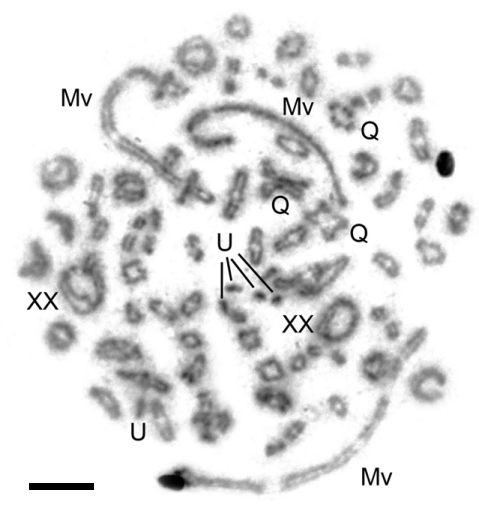

**Figure S1.** Tetraploid spermatocyte of *E. tancrei* specimen, # 25187, 2n=49, the diakinesis stage. Quadrivalents were formed by homologous acrocentrics (Q), multivalents were originated due to heterozygosity for Robertsonian translocations (Mv), univalents marked as U, two sex bivalents marked as XX. Scale bar = 5 µm.

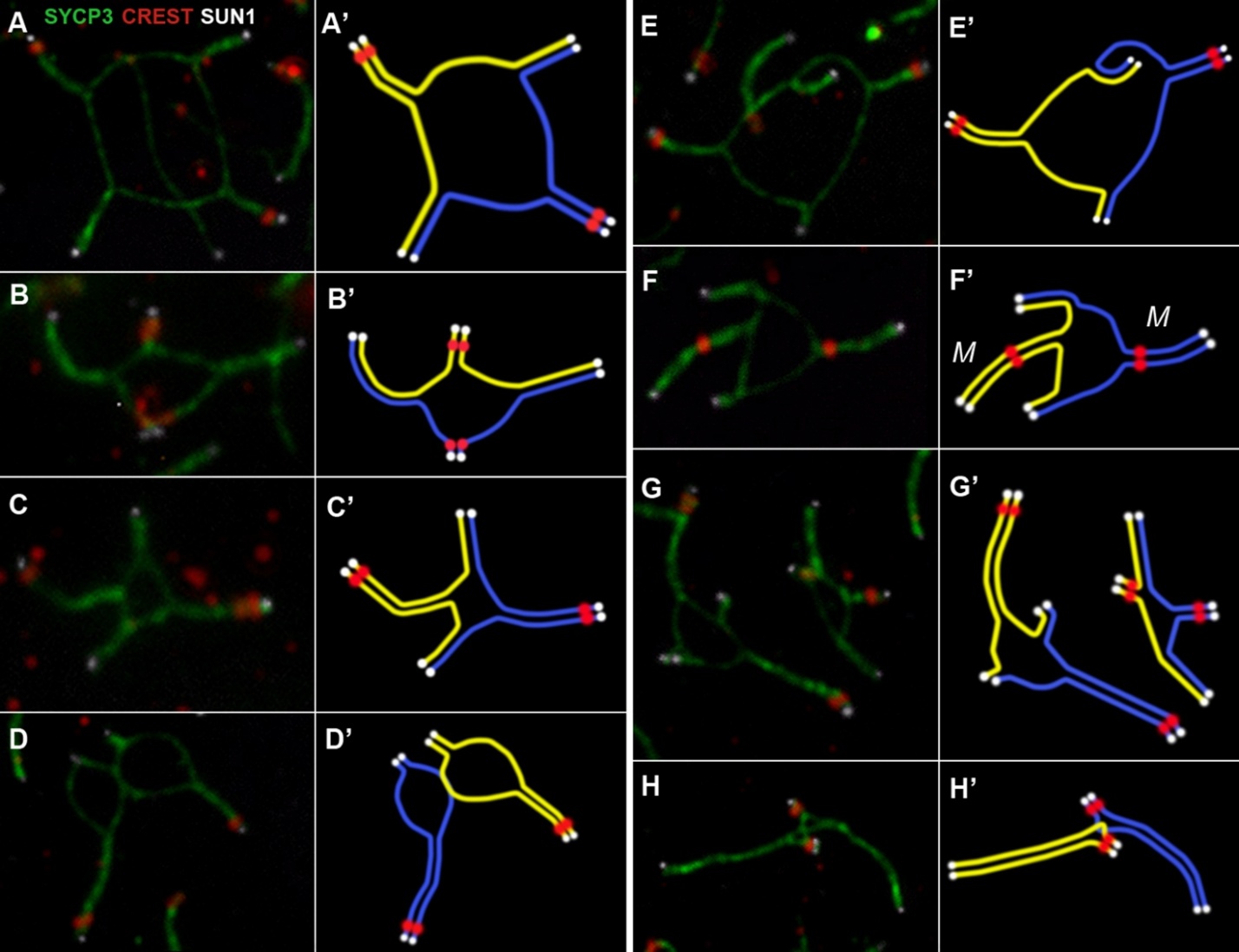

**Figure S2.** Various configurations of chromosomal quadrivalents in tetraploid spermatocytes of *E. alaicus* (A-H). Axial and lateral elements of the SCs were identified using an anti-SYCP3 antibody (green). Centromeres were identified using "CREST"—antibodies to kinetochore proteins (red). LINСs in the nuclear envelope were identified using SUN1 antibodies (white). All quadrivalents were taken from two meiotic nuclei presented in Supplementary Fig. 1. Each quadrivalent has an interpretation scheme (A’-H’). In panels F', 'M' labels metacentric chromosome configurations.

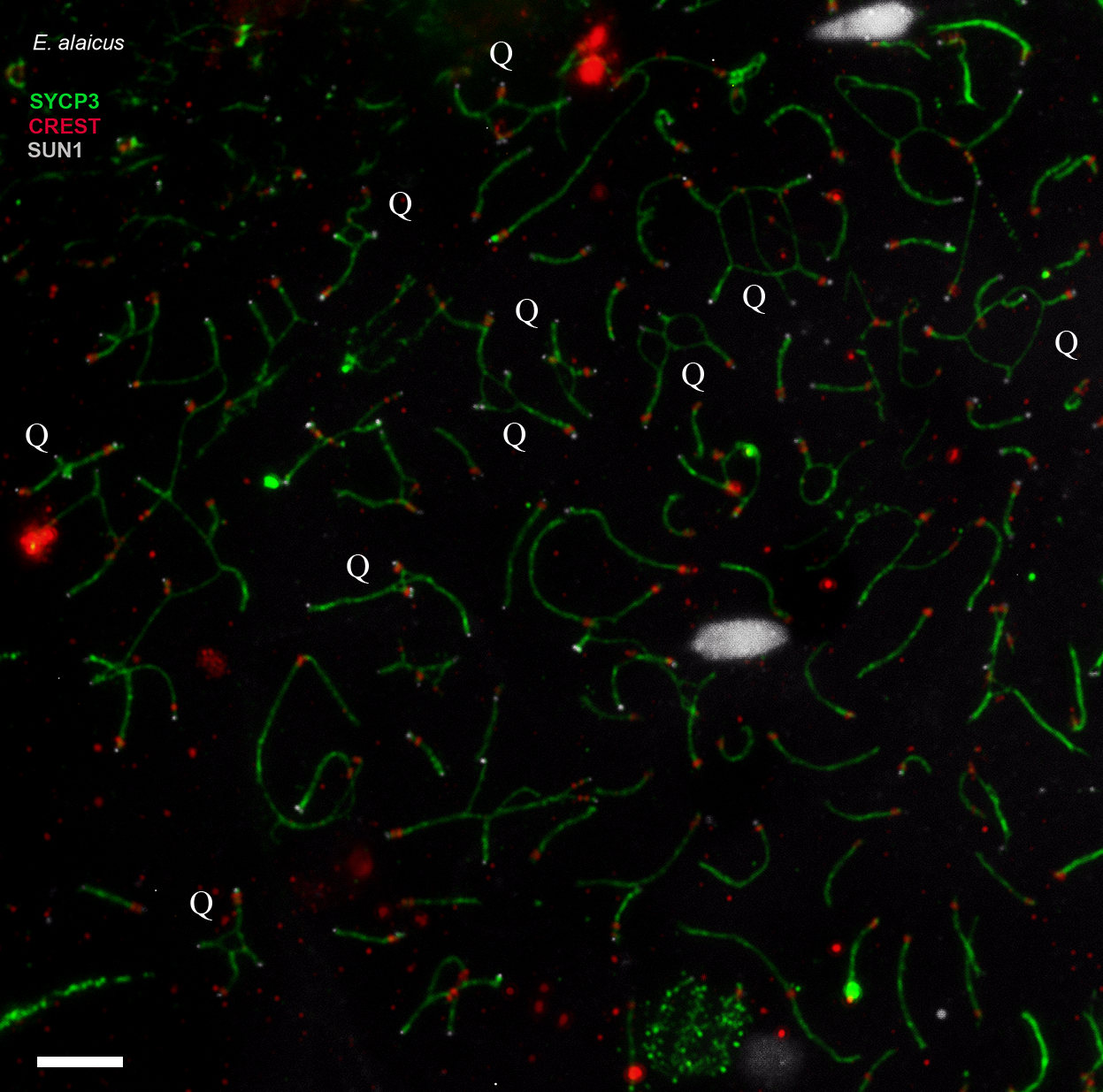

**Figure S3.** Two tetraploid and several diploid spermatocytes of *E. alaicus,* the pachytene stage. Axial and lateral elements of the SCs were identified using an anti-SYCP3 antibody (green). Centromeres were identified using "CREST"—antibodies to kinetochore proteins (red). LINСs in the nuclear envelope were identified using SUN1 antibodies (white). Q – chromosomal quadrivalent. Scale bar = 5 µm.

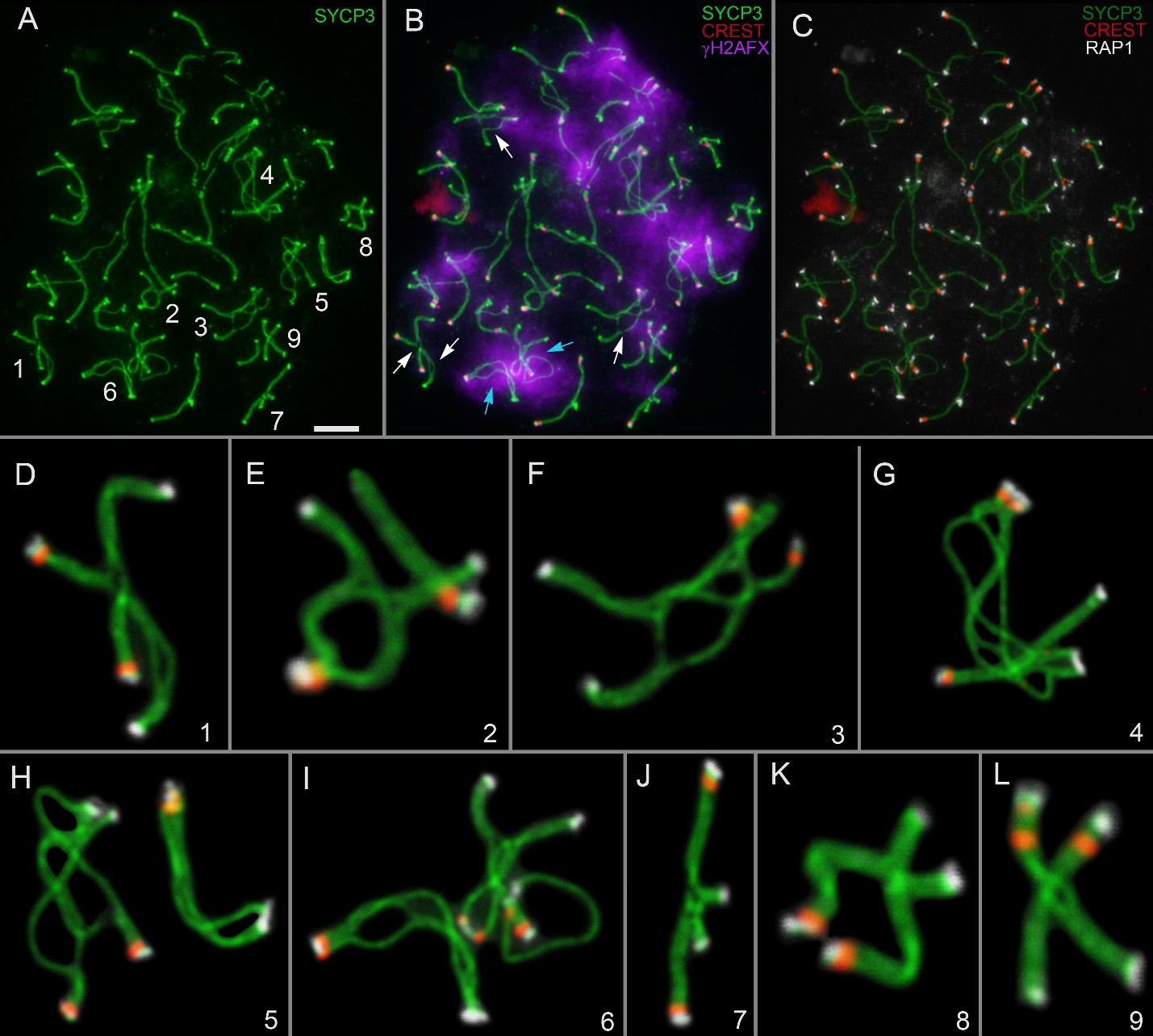

**Figure S4.** Tetraploid pachytene spermatocytes of E. talpinus (A-C). Axial and lateral elements of the SCs were identified using an anti-SYCP3 antibody (green); centromeres were identified using "CREST"—antibodies to kinetochore proteins (red); telomeres were detected using anti-RAP1 antibody (white); and chromatin inactivation was revealed using an anti-γH2AFX antibody (violet). Enlarged quadrivalents are shown in panels D-L. Cyan arrows indicate γH2AFX-positive asynaptic areas. White arrows indicate γH2AFX-negative chromosomal regions. Scale bar = 5 µm.

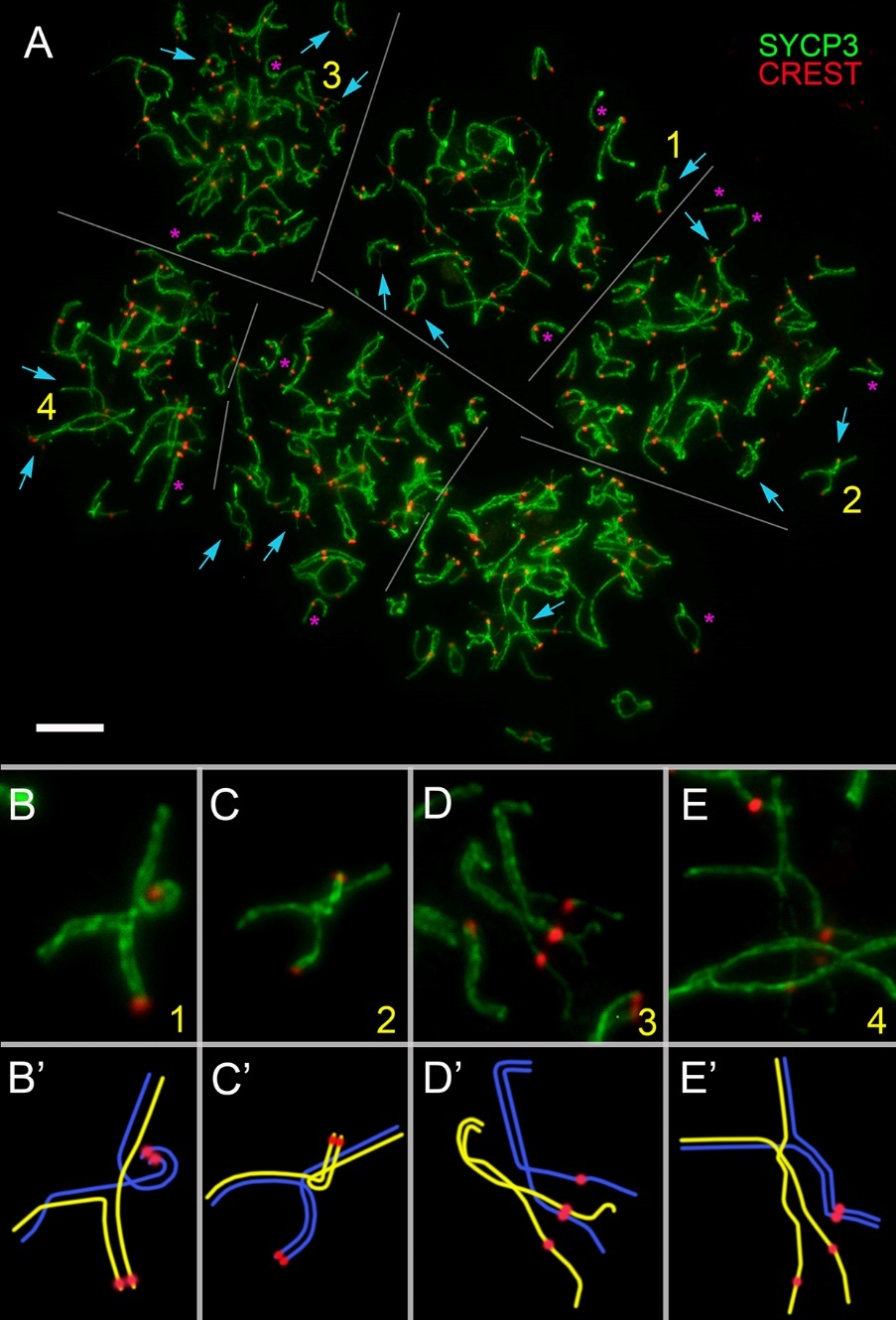

**Figure S5.** Six tetraploid pachytene spermatocytes of the mole rat N. leucodon (A). Axial and lateral elements of the SCs were identified using the anti-SYCP3 antibodies (green); centromeres were identified using "CREST"—antibodies to kinetochore proteins (white). Yellow numbers indicate those parts of the nuclei that are enlarged in insets (B-E). Cyan arrows point to SC quadrivalents and quadrivalent-like SC configurations. Magenta asterisks indicate separate SC bivalents. Scale bar (A) = 5 µm.
